# Social partner cooperativeness influences brain *oxytocin* transcription in Trinidadian guppies (*Poecilia reticulata*)

**DOI:** 10.1101/2021.03.01.433346

**Authors:** Sylvia Dimitriadou, Eduarda M. Santos, Darren P. Croft, Ronny van Aerle, Indar W. Ramnarine, Amy L. Filby, Safi K. Darden

**Affiliations:** Centre for Research in Animal Behaviour, University of Exeter, Exeter, UK; Biosciences, College of Life and Environmental Sciences, University of Exeter, Exeter, UK; Sustainable Aquaculture Futures, Biosciences, College of Life and Environmental Sciences, University of Exeter, Exeter, UK; International Centre of Excellence for Aquatic Animal Health, Cefas Weymouth Laboratory, Weymouth, UK; Department of Life Sciences, University of West Indies, St. Augustine, Trinidad and Tobago

**Keywords:** Cooperation, nonapeptides, isotocin, oxytocin, gene transcription, teleost

## Abstract

For non-kin cooperation to be maintained, individuals need to respond adaptively to the cooperative behaviour of their social partners. Currently, however, little is known about the biological responses of individuals to experiencing cooperation. Here, we quantify the neuroregulatory response of Trinidadian guppies (*Poecilia reticulata*) experiencing cooperation or defection by examining the transcriptional response of the *oxytocin* gene (*oxt*; also known as *isotocin*), which has been implicated in cooperative decision-making. We exposed wild-caught females to social environments where partners either cooperated or defected during predator inspection, or to a control (non-predator inspection) context, and quantified the relative transcription of the *oxt* gene. We tested an experimental group, originating from a site where individuals are under high predation threat and have previous experience of large aquatic predators (HP), and a control group, where individuals are under low predation threat and naïve to large aquatic predators (LP). In HP, but not LP, fish brain mid-section *oxt* relative transcription varied depending on social partner behaviour. HP fish experiencing cooperation during predator inspection had lower *oxt* transcription than those experiencing defection. This effect was not present in the control population or in the control context, where the behaviour of social partners did not affect *oxt* transcription. Our findings provide insight into the neuromodulation underpinning behavioural responses to social experiences, and ultimately to the proximate mechanisms underlying social decision-making.

## Introduction

Across taxa, individuals often exhibit cooperative behaviour, incurring costs to provide benefits to others [1–3]. Cooperative behaviour has posed a conundrum for the theory of natural selection when beneficiaries are non-kin [e.g. 2,4,5]. Theoretical work on cooperative behaviour has extensively examined the behavioural rules and potential strategies that ensure that cooperating individuals ultimately receive some reciprocation of benefits, leading to the stabilisation of cooperation in a population [5–8]. Unravelling the mechanisms underlying behavioural rules and strategies that underpin cooperation requires an understanding of the biological responses of individual appraisal of the cooperative behaviour of others. Although empirical studies have made progress in understanding the biological drivers underlying the performance of a cooperative act, there has been much less progress in documenting the neuromodulatory mechanisms underlying how individuals experience the cooperative behaviour of others.

Nonapeptide hormones in the brain are a likely key class of regulators that contribute to social decision-making in cooperative contexts, as they are amongst the most important modulators of social processes and emotions or emotion-like states across phylogenetic groups [9–11]. Oxytocin (OXT) in particular has been a focus of study in human empirical work on cooperation [12–15], and its teleost orthologue, oxytocin (Oxt; also known as isotocin – from here and onwards we use the fish nomenclature for Oxt) (https://zfin.org/ZDB-GENE-030407-1), has been implicated in cooperative interactions between labrid cleaner fish and their clients [16,17]. Overall, OXT is thought to facilitate social encounters by linking them to experiencing reward [18–20]. Although relatively little is known about the response of nonapeptide systems to experiencing cooperation or defection from the social environment, OXT has been shown to downregulate fear of social betrayal [15,21,22], and, in the presence of prior information about social partners, increase trust in humans [18,22]. Research also suggests that OXT inoculates the effects of experiencing a breach of trust in humans [13,21]. We might thus expect that OXT and its orthologues could be important mediators of how individuals appraise their environment as a function of the cooperativeness of social partners.

Here, we use female Trinidadian guppies (*Poecilia reticulata*) to explore the neuroregulatory response of individuals experiencing ostensibly cooperating and defecting social partners, by examining transcription of the *oxt* gene in the brain. Social experiences have been shown to affect gene transcription in teleosts [e.g. 23,24]; these responses can be both large (in terms of the number of genes affected) and fast (within 30 minutes from the presentation of the stimulus) [25]. Guppies cooperate in the context of predator inspection, a behaviour in which, in the presence of a potential predator, a single individual or a small number of fish leave the relative safety of the shoal or other refuge to approach and assess the potential threat; inspecting fish return to the shoal after collecting information about the level of the threat, and transmit this information to the remainder of the shoal [26–28]. Given that all shoal members benefit from the information collected by inspectors irrespective of whether they performed an inspection themselves [29–31], predator inspection is a good model for the study of cooperative behaviour [29]. Inspecting individuals are thought to be subjected to increased predation risk [30,32], which is shared when inspecting fish form cooperative partnerships [29,33]. Using this study system, we explore the effects of experiencing cooperation or defection from the social environment during predator inspection on the neuroregulatory response of the test fish. Given the well documented involvement of the oxytocin system in prosocial behaviour, we hypothesise that social experiences of cooperation from conspecifics will elicit greater relative transcription of the *oxt* gene than experiencing defection in this context.

## Methods

### Study subjects

Trinidadian guppies were collected from two populations, one living under risk of predation by large pisciverous fish (HP) (lower Guanapo) (10°36’31.5”N, 61°15’49.7”W) and one living without risk of predation by large pisciverous fish (LP) (upper Guanapo) (10°42’06.1”N 61°16’54.4”W) site of the Guanapo River on the island of Trinidad, and were transferred into housing tanks at the facilities of the University of West Indies (St. Augustine, Trinidad and Tobago). We chose to assay not only the HP fish (experimental group), but also the LP fish to act as a control group. LP habitats are characterised by the absence of major guppy predators such as pike cichlids (*Crenicichla frenata*) or *Hoplias malabaricus* [34,35]; this lack of predatory fish in LP habitats suggests that predator inspection is less important for survival in LP populations [35]. We would therefore not expect these fish to exhibit a biological response mediated by previous social experiences or strong selective pressure in the inspection context. While there is some gene flow between upstream and downstream populations, this is limited by the presence of waterfalls, and is primarily unidirectionally downstream [36,37]. The fish were fed with commercial flake once a day and were kept in a constant room temperature of 25°C, with a 12 hour light: 12 hour dark cycle. Thirty-four adult female guppies from each population were tested; this sample size was defined based on pilot data.

All experimental protocols were approved by the U.K. Home Office (Animal Scientific Licensing), and experiments were undertaken under a U.K. Home Office project licence (30/3430) and personal licence (I002BDF3F). All methods complied with the ARRIVE guidelines and were performed in accordance with the Animals (Scientific Procedures) Act 1986 (ASPA).

### Experimental paradigm

Fish were subjected to a predator inspection paradigm based on those previously used for this species [32,38,39]. The aim was to measure brain transcription of *oxt* in focal individuals when inspecting a predator in the presence of shoaling partners where either a shoal member ostensibly formed a cooperative partnership with the focal (joined the focal during inspection; here and throughout ‘cooperation’) or all shoal members ostensibly defected (did not join during inspection; here and throughout ‘defection’). As such, the experimental condition (predator) manipulated whether focal individuals ostensibly experienced their social partners as cooperating or defecting during predator inspection. In addition to this experimental condition, we also had a control condition that aimed to highlight any confounds of our experimental setup if they were present. In this control condition, the setup was analogous to the experimental condition, except that fish were not exposed to a predator that they could inspect (instead they had a familiar object) and as such did not have any experience of the cooperativeness of partners during inspection (i.e. the aim of our experimental condition), but ostensibly had an analogous experience of social partner presence when active in the observation lane (see below). As such, the control condition replicated all aspects of the experimental setup that may have affected gene transcription outside of the cooperative context (i.e. predator inspection), including the effects of having a mirrored or no-mirrored lane (such as the perceived size of the inspection lane). Each focal individual was assigned to either the experimental or the control condition.

The experimental arena consisted of two independent observation lanes separated by a Perspex barrier (Fig. 1). At the end of each observation lane, separated by a clear Perspex barrier, was the stimulus compartment, allowing for the transmission of visual cues only. The experimental or control stimulus was placed in this compartment. Individuals were either presented with a realistic model of *Crenicichla frenata*, a predator commonly found in high predation habitats (experimental condition) (*C. frenata* model: total length 12 cm) or a plastic aquarium plant (control condition). The use of predator models is well established in predator inspection studies [e.g. 27,40,41] as, contrary to live predators, they offer standardised predator behaviour, while still eliciting an anti-predator response. A small plastic plant was placed at the opposite end of the observation lane, providing a refuge for the focal fish. A small, clear, perforated cylinder containing the stimulus fish was also placed in the refuge area. Cooperative partnership during an inspection was simulated by placing a mirror along the observation lane so that focal individuals were inspecting alongside their mirror image, a behaviour that has been demonstrated to correlate with their behaviour during inspection with a live partner [39]. To simulate defection, focal individuals inspected alongside an opaque, non-reflective, surface. The mirror and no-mirror treatments were also used in the control condition to similarly simulate an accompanying fish (or no accompanying fish) when moving alongside the observation lane.

**Fig. 1.**
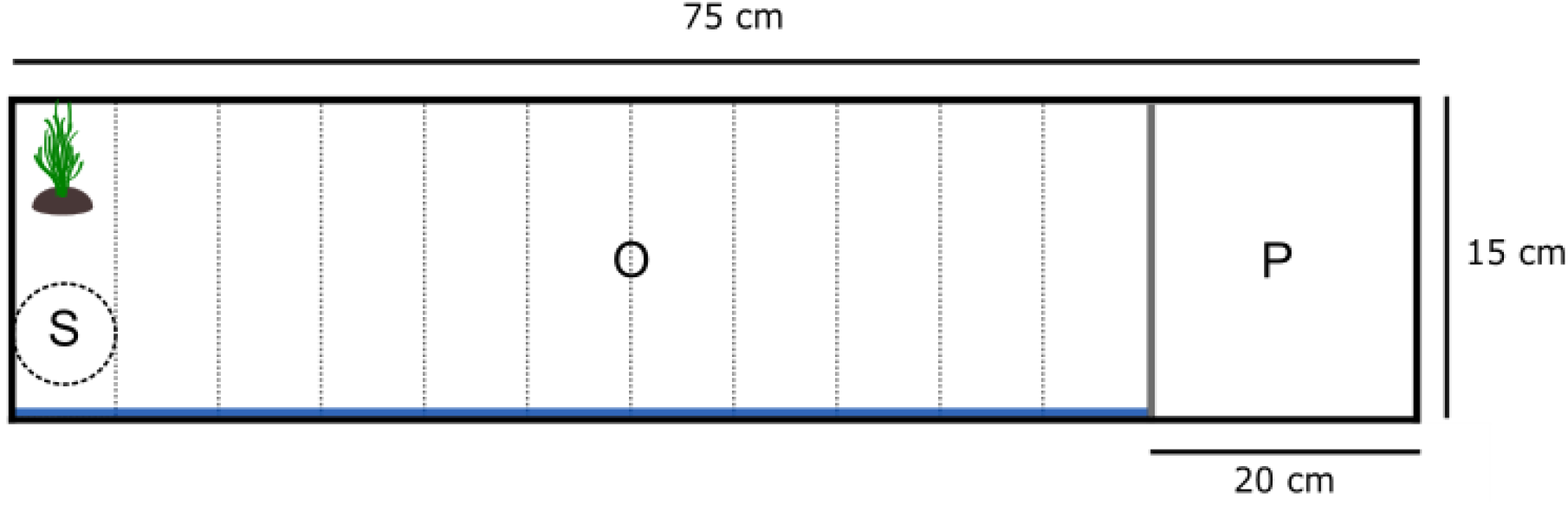
Schematic top view of the experimental set up for the predator inspection assay. A standard predator inspection tank (75×31×60 cm) with two observation lanes (O) divided by opaque Perspex was used. The predator model (or the plastic plant for the control condition) was placed in the predator compartment (P) behind a clear barrier that allowed for the transmission of visual cues. The stimulus shoal was placed in a clear, perforated cylinder (S) in the refuge area. Cooperation (partner) was simulated with the use of a mirror placed along the length of the inspection lane (blue line); defection (no partner) was simulated with the use of an opaque surface in lieu. The time the focal fish spent in each zone (vertical dotted lines) was recorded.

Three size-matched, same sex, stimulus fish drawn from a separate housing tank to the focal individual were introduced in the clear cylinder and left to acclimate for 10 minutes. At the end of this period, the focal individual was placed in the observation lane, with no visual access to the stimulus compartment (i.e. predator or plant stimulus hidden behind an opaque barrier), and was left for a further 10 minutes to acclimate. The focal fish had visual and olfactory access to the stimulus shoal throughout this period. Before the start of the trial, the focal fish was gently herded with a hand net to the refuge area, and the opaque barrier obstructing visual access to the stimulus compartment was lifted. The trial lasted for 5 minutes, after which visual access to the stimulus compartment was again obstructed. The focal individual remained in the observation lane for a further 10 minutes; it was then removed from the tank and rapidly euthanised using an overdose of tricaine methane sulfonate (MS 222, TMS; Sigma-Aldrich). The brain was removed and stored in RNA stabilisation reagent (RNA*later*; Qiagen) at −20°C. The samples were later dissected into three macro-areas [fore-section (telencephalon, habenula and preoptic area, excluding the olfactory bulbs and the hypothalamus), mid-section (including the optic tectum, diencephalon, and the hypothalamus) and hind-section (cerebellum and medulla oblongata)] and flash-frozen at −80°C (see Supplementary material). The mid-section samples were then used for the molecular analysis of the transcription of *oxt*.

Inspection trials were video recorded and the spatial position of focal fish within observation lanes was analysed with the Observer XT software (Noldus, the Netherlands) by a single observer blind to the population sample and condition. Inspection lanes were divided into 11 equidistant zones (5 cm length per zone), and the time spent in each zone was recorded. The average zone that fish occupied during inspection trials was then calculated. Eleven videos were excluded because of issues with the recording equipment.

### Analysis of transcription by qualitative real-time PCR

Total RNA from 57 fish was isolated from the excised tissues using the RNeasy Micro Kit (Qiagen), following the manufacturer’s instructions, including on-column treatment with RNase-free DNase (Qiagen). To estimate total RNA concentration, absorbance at 260 nm was measured (NanoDrop 1000, Thermo Fischer Scientific, Wilmington, DE, USA). RNA quality was verified by A_260nm_/A_280nm_ ratios; only samples with ratios ≥1.8 were used in downstream analysis (n=5–7 per experimental group). We present only the results for mid-section, as it was not possible to obtain RNA of sufficient quantity for the other two sections. RNA was then reverse transcribed using Moloney-Murine Leukaemia Virus (M-MLV) reverse transcriptase (Promega) according to the manufacturer’s instructions, using random hexamers (Eurofins Genomics) in a 96-well PCR machine (MasterCycler, Eppendorf, Mississauga, ON, Canada).

The cDNA sequences of *oxytocin* (*oxt*) and *ribosomal protein L8* (*rpl8*) were retrieved from the Ensembl Guppy database [Guppy_female_1.0_MT; 42] and specific primers were designed such that they spanned an intron, and ensuring that the primer pair would amplify all known transcripts, using Beacon Designer 3.0 (Premier Biosoft International, Palo Alto, CA, USA) and purchased from Eurofins Genomics. The following primer pairs were used: *oxt-FP*: 5’-GGTGGGAGAGCCTGTGG-3’/*oxt-RP*: 5’-GGTTCGGTGAGAAGTGTGG-3’ and *rpl8-FP*: 5’-GGAAAGGTGCTGCTAAACTC-3’/*rpl8-RP*: 5’-GGGTCGTGGATGATGTC-3’, resulting in amplicons of 198bp and 88bp in length, respectively. Annealing temperatures for the primers for the *oxt* and *rpl8* genes were optimised using a temperature gradient programme. The linearity and real-time PCR amplification efficiency (E; E=10^(−1/slope)^) of each pair of primers was calculated by running real-time PCR amplifications on a 10-fold dilution series of guppy midbrain cDNA from pooled, randomly selected samples, according to previously established protocols [43].

Real-time quantitative PCR using SYBR green chemistry was performed using the i-Cycler iQ Real-time Detection System (Bio-Rad Laboratories, Inc., Hercules, CA, USA). Samples were amplified in triplicate in 96-well optical plates (Fischer Scientific) in a 15 μl reaction volume, containing 1.5 μl cDNA, 7.5 μl iTaq Universal SYBR green mix (Bio-Rad), 5.25 μl of HPLC-grade water (Fisher Scientific) and 0.375 μl of each appropriate primer. *Taq* polymerase was activated by an initial denaturation step at 95°C for 15 minutes, followed by 45 cycles of denaturation at 95°C for 10 seconds and annealing at 58°C (*rpl8*) or 60°C (*oxt*) for 45 seconds, followed by melt curve analysis. Template-minus negative controls were run for each plate. A guppy midbrain cDNA mix of three random samples was repeatedly quantified on each plate to account for intra- and inter-assay variability (this positive control was run in duplicate on all plates – the average Ct differed ±0.3 among plates). To quantify differences in transcription between samples, *oxt* transcription values were normalised to *rpl8*, a housekeeping gene found to be consistently expressed in brain tissue in previous studies [44,45]. Relative transcription of the *oxt* was calculated using the arithmetic comparative method (2^-ΔΔCt^) as described by Filby et al. [44], which corrects for differences in PCR amplification efficiency between the target and housekeeping gene [46] (the efficiency for *oxt* was 2.170, and for *rpl8* was 2.028; in both cases the correlation coefficient for the standard curves was >0.994). Results were expressed as relative transcription ratios. The experimenter was blind to the experimental group of the samples during the analysis of gene transcription.

### Statistical analysis

All analyses were carried out in R v3.5.2 [47]. The average zone occupied during the behavioural trial was analysed using linear models. The global model included Condition (Control/Predator), Population (HP/LP) and Social environment (Partner/No partner), as well as the three-way interaction between these factors. Non-significant interactions were removed. The final model included Condition and the two-way interaction between Population and Social environment. Pairwise contrasts were obtained using the ‘lsmeans’ package v.2.30-0 [48], after multivariate *t* adjustment. Mid-section *oxt* transcription was analysed using beta regression in the ‘betareg’ v.3.0-5 R package [49]. Beta regression allows for statistical modelling of continuous data restricted to the unit interval (0,1) to model percentages and proportions [50,51]. The global model included Condition (Control/Predator), Population (HP/LP), and Social environment (Partner/No partner), and the three-way interaction between those factors. The average zone occupied in the observation lane during a trial was also included to control for any effects of overall proximity of the focal individual to the stimulus shoal. Comparisons of estimated marginal means (EM means) with false discovery rate corrections were carried out using the ‘emmeans’ v1.3.2 R package [52] separately for HP and LP populations. Power analysis was conducted using the ‘pwr’ R package v1.3-0 [53].

## Results

The average zone of the observation lane that individuals occupied during a trial was significantly affected by the interaction between Population and Social environment (Fig. 2) (F=6.1104, p=0.0182). *Post hoc* comparisons showed that, whereas the mean position during the behavioural trials was not affected by the Social environment in HP females, LP females without a partner (Social environment = no partner) did not approach the stimulus as closely as LP females with a partner, irrespective of the type of stimulus (predator vs. plant) (t-ratio=2.790, p=0.0396).

**Fig. 2.**
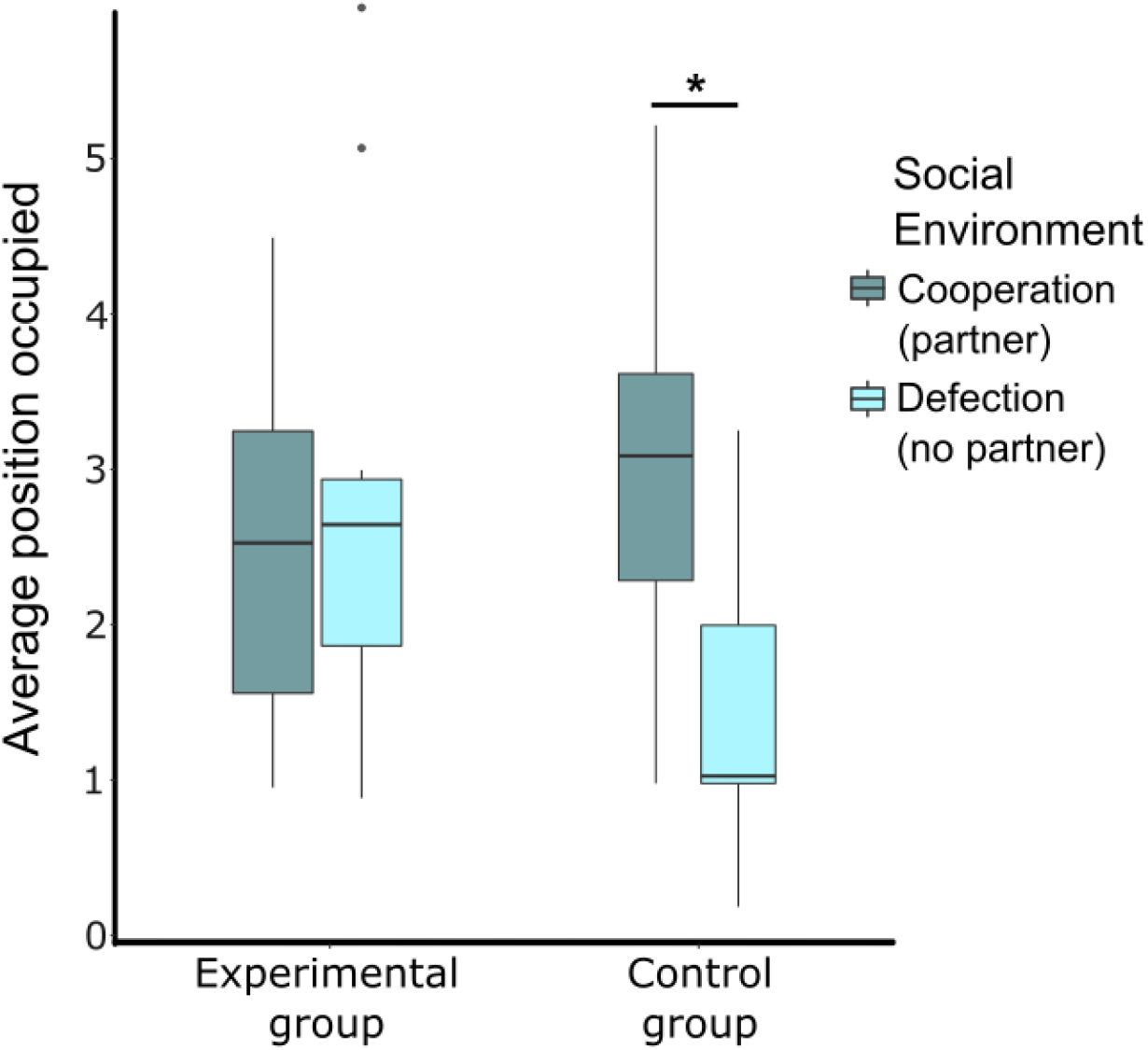
Spatial positioning of focal fish in the observation lane across treatments for experimental (originating from a population living under high predation risk, HP) and control (originating from a population living under low predation risk, LP) groups (n=5– 7 per group). The boxes represent the interquartile range (25th and 75th quartiles), and the horizontal lines represent the medians. The upper whisker extends to the largest value no further than 1.5 times the interquartile range (1.5*IQR), while the lower whisker extends to the smallest value within 1.5 times the interquartile range (Tukey boxplot). The dots represent outlying values. Females in the control group occupied areas of the observation lane further from the stimulus shoal and closer to the ‘predator’ compartment (containing either a plant or predator stimulus) (i.e. a higher zone value in the graph) in the presence of a simulated partner compared with those without a simulated partner. *: p< 0.05

We found a significant three-way interaction between Condition, Population and Social environment on *oxt* relative expression in the mid-section (Fig. 3) (‘Condition *Population*Social environment’: χ^2^(8, 41)=−2.09815, p=0.0146) (Pseudo R-squared=0.4795). Comparisons of estimated marginal means showed that whereas there was no difference in *oxt* relative transcription between experimental groups in LP fish, HP fish differed depending on the interaction between Condition (Experimental vs. Control) and the Social environment (Partner vs. No partner) (Fig. 3, left). Specifically, HP females inspecting a predator whilst experiencing cooperation (‘Partner’) had 9.3*10^-2^-fold lower *oxt* relative transcription than those experiencing defection (‘No partner’) (z-ratio=-3.038, p=0.005). The opposite effect was found in individuals in the control (plant) treatment, where those with a partner had 1.3*10^−1^-fold higher *oxt* transcription than those without a partner (z-ratio=5.219, p<0.001). Furthermore, individuals experiencing defection during predator inspection had 1.5*10^−1^-fold higher *oxt* relative transcription than those with the corresponding Social environment in the Control (plant) condition (z-ratio=-5.506, p<0.001). Relative *oxt* transcription in HP females experiencing cooperation did not differ between the experimental (predator) and control (plant) conditions (z-ratio=-2.010, p=0.067), suggesting that the interaction effect was driven by how *oxt* relative transcription was affected by a social environment where individuals chose to defect during predator inspection. The average zone of the observation lane used by an individual during a trial did not affect *oxt* transcription.

**Fig. 3.**
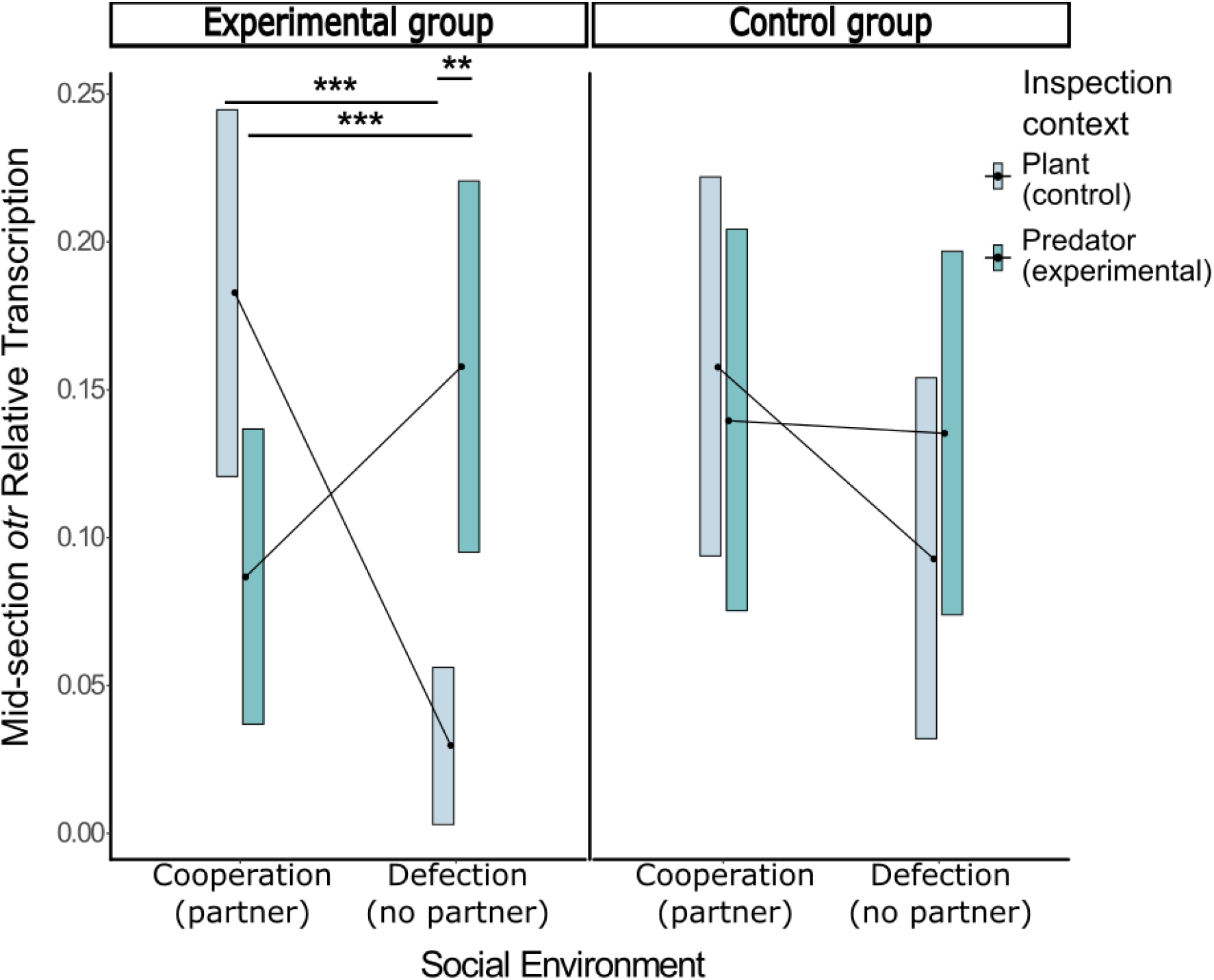
Mid-section *oxytocin (oxt)* transcription relative to *ribosomal protein L8 (rpl8)* transcription for females fish in the experimental group (originating from a population living under high predation risk, HP) (left) (n=23) and the control group (originating from a population living under low predation risk, HP) (right) (n=22) populations (n=5-7 per group). Lines represent estimated marginal means from beta regression, and boxes represent 95% confidence intervals. Female guppies originating from a High Predation habitat (Left) showed differential *oxt* transcription depending on the inspection context and cooperative environment. We found no such effect in guppies originating from a Low Predation habitat (Right). ** p< 0.01; *** p <0.001.

## Discussion

The aim of this study was to investigate the neuroregulatory response associated with experiencing cooperation or defection during predator inspection by examining brain gene transcription for *oxt*. We found that in fish originating from a HP habitat, *oxt* transcription in the brain mid-section varied as a function of inspecting a predator in a partnership as opposed to singly when social partners were available to join. This was not the case for control fish originating from a LP habitat, where *oxt* transcription was found to be independent of the behaviour of social partners during predator inspection. To our knowledge, this is the first study to examine the effects of the experience of social partner behaviour in a cooperative context on brain nonapeptide transcription patterns. Our study provides insight into the neuromodulatory mechanisms underlying cooperative strategies and behavioural rules associated with subsequent decision-making in such contexts.

The response of nonapeptide systems to experiences of cooperation or defection from the social environment remains relatively unexplored; however, there is evidence demonstrating a role for OXT and its orthologues in the expression of cooperative behaviour. For example, intranasal OXT administration has been shown to increase cooperative behaviour in humans playing an iterated Prisoner’s Dilemma game [13]. Also, Cardoso and colleagues [17] demonstrated a sex-dependent relationship between Oxt levels in the forebrain of *Labroides dimidiatus* and honesty during cooperative interactions with client reef fish, thus implying a role of the Oxt pathways in mutualistic behaviour. Furthermore, in this same study system, client presence results in elevated forebrain Oxt levels, suggesting a link between environmental appraisal in a cooperative context and the forebrain Oxt pathway [16]. Our results are consistent with the documented involvement of Oxt systems in cooperative contexts, and imply their involvement in the response to cooperation or defection from an individual’s social environment; however, contrary to our predictions, experiencing defection from the social environment during inspection led to higher *oxt* transcription compared with experiencing cooperation (and compared to our control) in HP fish. It is not immediately clear why experiencing defection from social partners during predator inspection would lead to higher *oxt* transcription than cooperation, but findings reported for humans may inform on constructing hypotheses to explain these results. In humans, OXT has been implicated in the inoculation of social betrayal [15,21,22]; it is possible that the effects of experiencing defection on *oxt* transcription observed here are reflecting a similar role of nonapeptide systems during negative (i.e. defection) social interactions. Given the proposed role of Oxt in the regulation of the salience of social stimuli in a context-dependent manner via the regulation of attention to social cues [54], it is also possible that the observed *oxt* transcription patterns represent a role the higher propensity for individuals to remember characteristics associated with cheaters compared to cooperators, a bias that has for example been demonstrated in humans [55]. This could be explained by the possibility that information regarding ‘cheating’ is more important when predicting trait characteristics of social partners, and therefore their future behaviour [56,57]; that is ‘negative’ cues can potentially be more diagnostic than ‘positive’ ones [57,58]. Indeed the most recent work in this species would suggest that female guppies are most socially selective following social partner defection compared to cooperation [58]. More work is needed to gain a better understanding of both the role of Oxt in behaviour in this cooperative context and in subsequent decision-making.

Social behaviour in vertebrates is largely modulated by a number of reciprocally connected nodes in the brain known as the ‘Social Behaviour Network’ (SBN) [59]. The SBN, alongside the nuclei forming the basal forebrain reward system constitute the ‘Social Decision-Making Network’ (SDMN) [60,61], which is thought to be the neural substrate of the regulation of social behaviours such as the ones considered paramount components of cooperative behaviour [62–64]. Several of the nodes of the SDMN, including the periaqueductal grey (PAG), the ventral tuberal nucleus (vTn – the teleostean homologue of the anterior hypothalamus), the anterial tuberal nucleus (aTn – homologous to the ventromedial hypothalamus) and the posterium tuberculum (TPp – homologous to the ventral tegmental area) are located within the mid-section of the brain analysed [62,65] and it is therefore perhaps not surprising that social cooperativeness affected the activity of Oxt systems in this section. This study, however, did not aim to detect brain nodes directly involved in the perception of cooperative behaviour. Further work is needed to identify the exact brain regions where *oxt* transcription is differentiated as a result of experiencing cooperation or defection from the social environment, using techniques that focus on detecting oxt activity in specific nodes of the SDMN, e.g. through immunohistology. Future work should also explore the extent to which the measured neuromodulatory patterns are subsequently linked with social decisions affecting the exhibition of specific strategies or rules [e.g. Walk-Away [8], generalized reciprocity [66], direct reciprocity [29]] through the manipulation of Oxt systems and an investigation of resulting cooperative and social decision-making.

Appraisal of the environmental context is expected to affect an individual’s internal state and therefore subsequent behaviour [64,66,67]. Contrary to fish in our experimental group (drawn from an HP population), *oxt* transcription in fish in our control group (drawn from an LP population) was not affected by social partner behaviour during predator inspection. Across rivers, LP habitats lack any of the major guppy predators such as pike cichlids [34,35] and predator inspection is thought to be less important for survival in these populations [35]. Our results were thus as expected for these two groups; any observed effect of social partner behaviour on *oxt* transcription would only manifest in our experimental group. If this is a pattern that occurs across guppy populations that differ in predation risk, then it could be evidence that the effect of the cooperativeness of social partners during predator inspection on subsequent individual decision-making is under differential selective pressure across populations [39]. Of course any difference could also be related to previous experience of cooperation and defection during inspection of a large pisciverous fish: the control group fish (LP) in our study were naïve to this type of predator and it is possible that repeated experiences with inspection of these types of predators in cooperative contexts would evoke a response of *oxt* transcription more akin to that observed in the experimental group fish (HP). Importantly, although our findings provide the first evidence to suggest that the response of neuromodulatory mechanisms to social experiences may differ among predation regimes in guppies, further work is needed to specifically test such a hypothesis that investigates whether the result is sustained within and across populations originating from different rivers.

Relative *oxt* transcription was not affected by an individual’s own behaviour in the observation lane. This would suggest that in our experimental condition where focal individuals were inspecting a predator, their own cooperativeness (propensity to approach the predator) was not related to *oxt* transcription at this time point. Indeed, the lack of an effect on relative transcription in this condition for fish in our control group (LP fish) where the presence and absence of a social partner affected spatial position in the observation lane supports this further. The lack of an effect however is somewhat unexpected as Oxt has been implicated in the expression of cooperative behaviour [16,17]: indeed there is a misalignment of difference in behaviour (LP fish) and difference in transcription (HP fish). It is possible that our experimental setup lacked the statistical power to show differences between the behaviour of HP fish under different experimental conditions; a power analysis showed an estimated statistical power of ~81%, suggesting that it is unlikely that we did not detect a true effect. We therefore believe that our results support the pattern that even if the expression of cooperative behaviour does not differ between HP individuals experiencing cooperation or defection, their response to such experiences, as expressed by the differences in *oxt* transcription differs. In order to fully understand the relationship between *oxt* gene transcription and cooperative behaviour during predator inspection, further work explicitly examining the effects of Oxt signalling on cooperation is needed.

It should also be noted that the effects on gene transcription shown here are extremely fast (15 minutes post trial start). As this study used only one sampling point, it is impossible to infer the timeline and magnitude of the response of Oxt. To our knowledge, this is the first study to look at the response of *oxt* transcription so shortly after a social experience, making it difficult to compare the effect found here to other studies; however, in teleosts, social stimuli have been shown to elicit genomic responses as fast as 30 minutes post stimulus exposure [68]. To further understand the effects of social experiences on the regulation of this gene, a time-course study is needed to investigate the temporal dynamics and amplitude of this effect.

Our findings demonstrate that in Trinidadian guppies originating from a HP locality, experiencing cooperation or defection from the social environment in the context of predator inspection affects *oxt* transcription patterns in the mid-section of the brain. Given the centrality of nonapeptide systems in the regulation of social behaviour, it is likely that the different *oxt* transcription profiles exhibited by HP fish in different experimental conditions will affect the focal individual’s subsequent social behaviour and decision-making, such as the formation of social bonds [69]. While more work is needed to explore whether experiencing cooperation or defection affects *oxt* transcription in other brain sections, and to identify the nodes underlying this effect, this study provides novel insight into the neuromodulation underpinning behavioural responses to social experiences, and furthers our understanding of the proximate mechanisms involved in social decision-making.

## List of abbreviations used

oxt: Oxytocin
HP: High Predation
LP: Low Predation
SBN: Social Behaviour Network
SDMN: Social Decision-Making Network
PAG: periaqueductal grey
vTn: ventral tuberal nucleus
aTn: anterial tuberal nucleus
TPp: posterium tuberculum

## Acknowledgements

The authors would like to thank Raj Mahabir and Rob Heathcote for assistance in the field, and Lisa Bickley and Hannah Littler for their help with the optimisation of the RNA extraction protocol.

## Authors contributions

S.D., S.K.D., D.P.C. and E.M.S. designed the study. R.v.A. and A.L.F. designed the primers. I.R. contributed materials and provided infrastructure while in Trinidad, and E.M.S. supervised the molecular analysis. S.D. carried out the behavioural experiments and molecular data collection, and S.K.D. analysed the behavioural trial videos. S.D. analysed the data, and in discussion with S.K.D. wrote the first draft of the manuscript. All authors contributed to the final version of the manuscript.

## Data availability statement

The raw data supporting this manuscript will be archived on Figshare upon acceptance.

