## Supplemental material for "Social partner cooperativeness influences brain *oxytocin* transcription in Trinidadian guppies (*Poecilia reticulata*)"

SUPPLEMENTARY TABLES

Table S1. Marginal effects of experimental condition, population and cooperative environment on average zone during inspection. Linear model with non-significant interactions removed. Statistically significant factors are shown in bold.

| | | Estimate | Std. error | t value | p value |
|---|---|---|---|---|---|
| **Intercept** | | **2.510** | **0.422** | **5.955** | **<0.001** |
| Inspection Context | Control (Plant) | 0 | - | - | - |
| | Predator | -0.160 | 0.372 | -0.430 | 0.670 |
| Population | HP | 0 | - | - | - |
| | LP | 0.586 | 0.523 | 1.120 | 0.270 |
| Cooperative Environment | Cooperation | 0 | - | - | - |
| | Defection | 0.363 | 0.523 | 0.695 | 0.492 |
| **Population * Cooperative Environment** | **HP – Cooperation** | **0** | **-** | **-** | **-** |
| | **LP – Defection** | **-1.830** | **0.740** | **-2.472** | **0.018** |

Table S2. Post hoc analysis for the ‘Population*Cooperative Environment’ interaction on average zone during inspection. Pairwise least squares means comparisons. Statistically significant contrasts are shown in bold.

| Contrast | Estimate | Std. error | t ratio | p value |
|---|---|---|---|---|
| HP/Cooperation - LP/Cooperation | -0.586 | 0.523 | -1.120 | 0.679 |
| HP/Cooperation - HP/Defection | -0.363 | 0.523 | -0.695 | 0.898 |
| HP/Cooperation - LP/Defection | 0.880 | 0.536 | 1.641 | 0.369 |
| LP/Cooperation - HP/Defection | 0.223 | 0.511 | 0.435 | 0.972 |
| **LP/Cooperation - LP/Defection** | **1.466** | **0.526** | **2.790** | **0.040** |
| HP/Defection - LP/Defection | 1.244 | 0.523 | 2.377 | 0.099 |

Table S3. Marginal effects of average zone during inspection, inspection context, population and cooperative environment on mid-section relative *oxt* expression. Beta regression model with no model selection. Statistically significant factors are shown in bold.

| | | Estimate | Std. error | z value | p value |
|---|---|---|---|---|---|
| **Intercept** | | **-1.695** | **0.313** | **-5.412** | **<0.001** |
| Average zone during inspection | | 0.080 | 0.089 | 0.902 | 0.367 |
| **Inspection context** | **Control (Plant)** | **0** | **-** | **-** | **-** |
| | **Predator** | **-0.857** | **0.381** | **-2.249** | **0.025** |
| Population | HP | 0 | - | - | - |
| | LP | -0.179 | 0.321 | -0.558 | 0.577 |

| | | Estimate | Std. error | z value | p value |
|---|---|---|---|---|---|
| **Cooperative Environment** | **Cooperation** | **0** | **-** | **-** | **-** |
| | **Defection** | **-1.986** | **0.507** | **-3.921** | **<0.001** |
| Exp. Condition * Population | Control – HP | 0 | - | - | - |
| | Predator – LP | 0.714 | 0.523 | 1.366 | 0.172 |
| **Exp. Condition * Coop. Environment** | **Control – Cooperation** | **0** | **-** | **-** | **-** |
| | **Predator – Defection** | **2.666** | **0.646** | **4.126** | **<0.001** |
| **Population * Coop. Environment** | **HP – Cooperation** | **0** | **-** | **-** | **-** |
| | **LP – Defection** | **1.382** | **0.690** | **2.005** | **0.045** |
| **Exp. Condition * Population * Coop. Environment** | **Control – HP – Cooperation** | **0** | **-** | **-** | **-** |
| | **Predator – LP – Defection** | **-2.098** | **0.859** | **-2.443** | **0.015** |

Table S4. Post hoc analysis for the ‘Inspection Context*Cooperative Environment’ interaction on midsection it expression in fish originating from the High Predation site. Pairwise estimated marginal means comparisons. Statistically significant contrasts are shown in bold.

| Contrast | Estimate | Std. error | z value | p value |
|---|---|---|---|---|
| Control/Cooperation - Predator/Cooperation | 0.0741 | 0.037 | 2.010 | 0.067 |
| **Control/Cooperation - Control/Defection** | **0.131** | **0.025** | **5.219** | **<0.001** |
| Control/Cooperation - Predator/Defection | 0.003 | 0.029 | 0.132 | 0.895 |
| Predator/Cooperation - Control/Defection | 0.0350 | 0.032 | 1.078 | 0.337 |
| **Predator/Cooperation - Predator/Defection** | **-0.093** | **0.031** | **-3.038** | **0.005** |
| **Control/Defection - Predator/Defection** | **-0.150** | **0.027** | **-5.506** | **<0.001** |

Table S5. Post hoc analysis for the ‘Inspection Context*Cooperative Environment’ interaction on midsection *oxt* expression in fish originating from the Low Predation site. Pairwise estimated marginal means comparisons. Statistically significant contrasts are shown in bold.

| Contrast | Estimate | Std. error | z value | p value |
|---|---|---|---|---|
| Control/Cooperation - Predator/Cooperation | -0.001 | 0.044 | -0.020 | 0.984 |
| Control/Cooperation - Control/Defection | 0.046 | 0.034 | 1.352 | 0.530 |
| Control/Cooperation - Predator/Defection | 0.003 | 0.024 | 0.143 | 0.984 |
| Predator/Cooperation - Control/Defection | 0.028 | 0.043 | 0.652 | 0.984 |
| Predator/Cooperation - Predator/Defection | -0.015 | 0.035 | -0.420 | 0.984 |
| Control/Defection - Predator/Defection | -0.061 | 0.043 | -1.444 | 0.529 |
